# Environmental Stochasticity Reshapes Persistence and Extinction Dynamics in a Fear-Mediated Two-Species Competitive System

**DOI:** 10.64898/2026.07.04.736416

**Authors:** Vaibhava Srivastava

## Abstract

Environmental variability can strongly alter coexistence among competing species and their extinction risk, particularly when population dynamics are shaped by behavioral interactions, such as fear. In this work, we develop a novel stochastic differential equation competition model that incorporates both non-consumptive fear effects and environmental variability to investigate how behavioral interactions influence species coexistence under random fluctuations. Our result reveals that environmental stochasticity can drive species to extinction even when the corresponding deterministic system admits coexistence. In particular, under an explicit stability condition on the fear and competition parameters and sufficiently strong averaged noise intensities, we prove that both competing species become extinct exponentially almost surely. Conversely, we derive a stochastic persistence criterion in terms of fear, competition, and noise-induced suppression parameters for the fearful species. We further demonstrate that environmental noise may reverse classical competition-exclusion outcomes, leading to qualitatively different long-term dynamics from those predicted deterministically. These results provide rigorous thresholds separating stochastic extinction from persistence and highlight the critical role of environmental variability in fear-mediated competitive ecosystems. From an applied perspective, these results provide insight into how behavioral interactions and environmental variability influence species survival, with potential applications in ecological management and conservation.

## Introduction

In an ecosystem, species compete for food, resources, and mates through competition. Competition between species has been extensively studied over the past few decades, often using the classical Lotka–Volterra competition model and its variants [1–3]. These models consider the growth of species as well as interspecific and intraspecific competition. They effectively predict important biological outcomes, such as coexistence, competitive exclusion, and bistability, and they have diverse applications in ecology and invasion science [4–14]. Let *x*_1_ and *x*_2_ represent the densities of the two competing species; their dynamics are described by this classical Lotka–Volterra competition model [15] as follows:

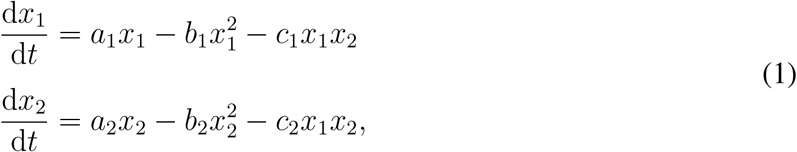

Here, *a*_1_ and *a*_2_ represent the intrinsic growth rates, while *b*_1_ and *b*_2_ represent the intraspecific competition rates. Additionally, *c*_1_ and *c*_2_ denote the interspecific competition rates for the respective species *x*_1_ and *x*_2_. All parameters considered above are positive.

Recently, in several ecological case studies, researchers observed that competitors also exert fear on each other. One notable case of fear among competing species is intraguild predation, a commonly observed ecological phenomenon in which potential competitors in the same “guild” (i.e., competing species) also prey upon each other. Such as non-consumptive effects exerted by intraguild predator mites (*Blattisocius dentriticus*) on their competitor (*Neoseiulus cucumeris*) [16]. There is also strong evidence that we have a fear effect among purely competing species. In the Pacific Northwest of the United States, the invasive species, the Barred owl, exerts fear (or co-occurrence effect) on the native northern spotted owls [17–21]. Barred owls (*Strix varia*) are a generalist species of owl, native to eastern North America. They have expanded their range westward over the last century and are considered invasive in western North America. Currently, their range overlaps with that of the spotted owl (*Strix occidentalis*), a specialist owl native to the northwest and western North America. Barred owls exert a negative influence on spotted owls due to several reasons, such as their large body size and exhibiting aggressive behavior towards spotted owls. Field observations report frequent attacks by barred owls on spotted owls, and even on surveyors who imitate spotted owl calls [21]. There is also evidence that barred owls aggressively chase spotted owls out of shared habitat, but not vice versa. Due to the presence of barred owls, spotted owls tend to show reduced responsiveness, which results in reduced territory defense, mate attraction, and lower fecundity. Recent ecological case studies and surveys suggest that barred owls will threaten spotted owls to their possible competitive exclusion [19].

There is also other recent empirical evidence to support the presence of fear among competing species. In a series of 6-year-long experiments in various Caribbean islands, researchers aim to refute the theory of adaptive predation. The theory suggests that predators reduce the dominance of dominant competitors, thereby preventing competitive exclusion and enhancing coexistence in food webs [22]. In their experiment, brown anolis (*Anolis sagrei*) and green anolis (*Anolis smaragdinus*) show that the introduction of the ground-dwelling curly-tailed lizard (*Leiocephalus carinatus*) induces fear in the brown anolis, prompting it to move up into the canopy where the green anolis resides. This results in intensified competition and a disruption and breakdown in co-existence. Interestingly, fecal analysis reveals that the curly-tailed lizard consumed only brown anolis in a small number of samples, suggesting that the brown anolis’ movement is primarily driven by fear rather than predation. Similar non-consumptive effects have been observed among competing aphid species.

The evidence strongly suggests the need to incorporate fear dynamics into a competitive two-species model in which one species is fearful of the other. In the recent work [23], authors formulate a classical two-species Lotka-Volterra competition model, where only one competitor instills fear in the other:

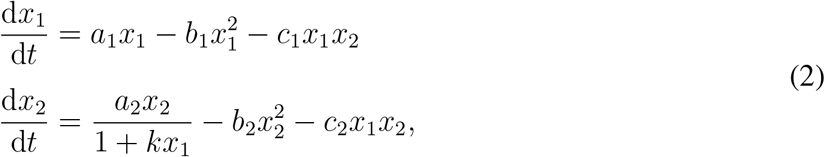

where *x*_1_ and *x*_2_ are the population densities of two competing species, *a*_1_ and *a*_2_ are the intrinsic (per capita) growth rates, *b*_1_ and *b*_2_ are the intraspecific competition rates, *c*_1_ and *c*_2_ are the interspecific competition rates, *k* is a fear coefficient. The term 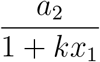 in the models denotes the reduction in the effective growth rate of species *x*_2_ due to the presence of species *x*_1_. Hence, as *x*_1_ increases, the growth of *x*_2_ is increasingly suppressed, which reflects behavioral or physiological responses to perceived risk. All parameters considered are positive. In [23], the authors found that fear can lead to several interesting dynamic effects in classical competitive scenarios, both in the constant (ordinary differential equation) and spatially heterogeneous (reaction-diffusion partial differential equation) cases. Notably, authors show novel bistability dynamics for certain levels of fear. All of these findings in both ecological and sociopolitical contexts relate to the concept of the “landscape of fear” (LOF) in ecology. There are other follow-up theoretical works in which researchers incorporated the fear mechanism to study the interplay between non-consumptive effects, such as fear, and other ecological mechanisms in various ecosystems [24, 25]. All of these studies clearly indicate that incorporating fear among competing species not only leads to a variety of novel dynamic behaviors but also aids in species management, thereby contributing to conservation biology.

While deterministic models (ODEs and PDEs) such as the above are useful for capturing population dynamics, they often overlook the random perturbations. These random perturbations can significantly impact long-term outcomes, such as extinction or coexistence. There are several ecological case studies that show that factors such as climatic variability, which includes fluctuations in temperature, precipitation, snow cover, and extreme events, can alter survival, reproduction, carrying capacity, and the strength of species interactions [26–31]. All of this variability can shape long-term population dynamics, affect species population growth and persistence, and drive species to extinction. For instance, a study by Vázquez et al. [26] showed that changes in the variance of vital rates can either increase or decrease stochastic population growth. Furthermore, the dynamics depend on factors such as life history, correlations among vital rates, and the temporal structure of environmental fluctuations.

There are also several ecological case studies that highlight the significance of stochasticity in population dynamics. A notable example is the interaction between the snowshoe hare (*Lepus americanus*) and the Canada lynx (*Lynx canadensis*) in the boreal forests of North America. A well-established result for this case study is that the system exhibits the well-known 9-11 year population cycles [27, 28]. For this case-study, Stenseth et al. [27] demonstrated that the hare dynamics are better described by a three-trophic-level structure involving vegetation, hares, and a predator guild, whereas lynx dynamics are effectively governed by a lower-dimensional hare-lynx interaction. Their analysis showed that deterministic skeleton models generate only damped oscillations rather than sustained cycles. They further demonstrated that environmental stochasticity can maintain these damped oscillations and create the persistent cyclic behavior observed in long-term field data. This finding emphasizes that stochasticity is not just a perturbation around deterministic dynamics, but rather an essential ecological mechanism for maintaining realistic population cycles. Later, Yan et al. studied and linked climate variability directly to the hare-lynx cycles using historical fur harvest records and generalized additive models [32]. Their study incorporated density dependence, asymmetric predation, and both global and local climatic variables, including the North Atlantic Oscillation (NAO), Southern Oscillation Index (SOI), Northern Hemisphere Temperature (NHT), rainfall, and snow cover. They found that density dependence and predator effects alone produced only damped oscillations, whereas the inclusion of climatic forcing generated persistent 10-year cycles that closely matched observed population fluctuations. There are also several empirical studies that suggest that the how changes in the habitat and climate can impact the population dynamics of Spotted Owls [33, 34].

These studies provide strong ecological justification for stochastic differential equation models. In systems involving fear effects, competition, and nonlinear trophic interactions, stochasticity may strengthen or weaken behavioral responses and shift the system away from deterministic predictions. Therefore, incorporating multiplicative environmental noise into the present competing-species model allows us to capture these realistic ecological uncertainties and better understand how random fluctuations and non-consumptive effects jointly shape long-term population dynamics. Therefore, incorporating stochasticity into the present competition model provides a more realistic framework for studying persistence, stability, and long-term species survival under environmental uncertainty. The stochastic formulation allows us to investigate how random environmental perturbations modify deterministic predictions, identify thresholds for persistence and extinction, and better understand the resilience of ecological systems subject to unpredictable disturbances [35–40].

To account for this environmental variability, we introduce multiplicative stochastic perturbations into the population dynamics. The resulting stochastic system is expressed as follows:

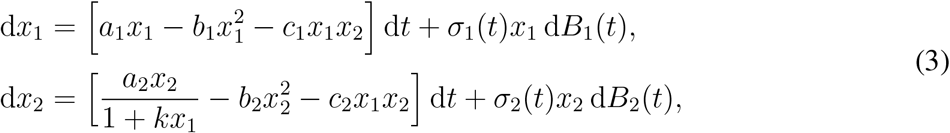

where *B*_1_(*t*) and *B*_2_(*t*) are statistically independent Brownian motions at time *t*, and *σ*_1_, *σ*_2_ ≥ 0 represent the intensity of environmental noise, and 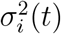 is a continuous bounded function on ℝ^+^. The multiplicative noise structure ensures that stochastic fluctuations scale with population size, which is consistent with ecological observations.

**Table 1:**
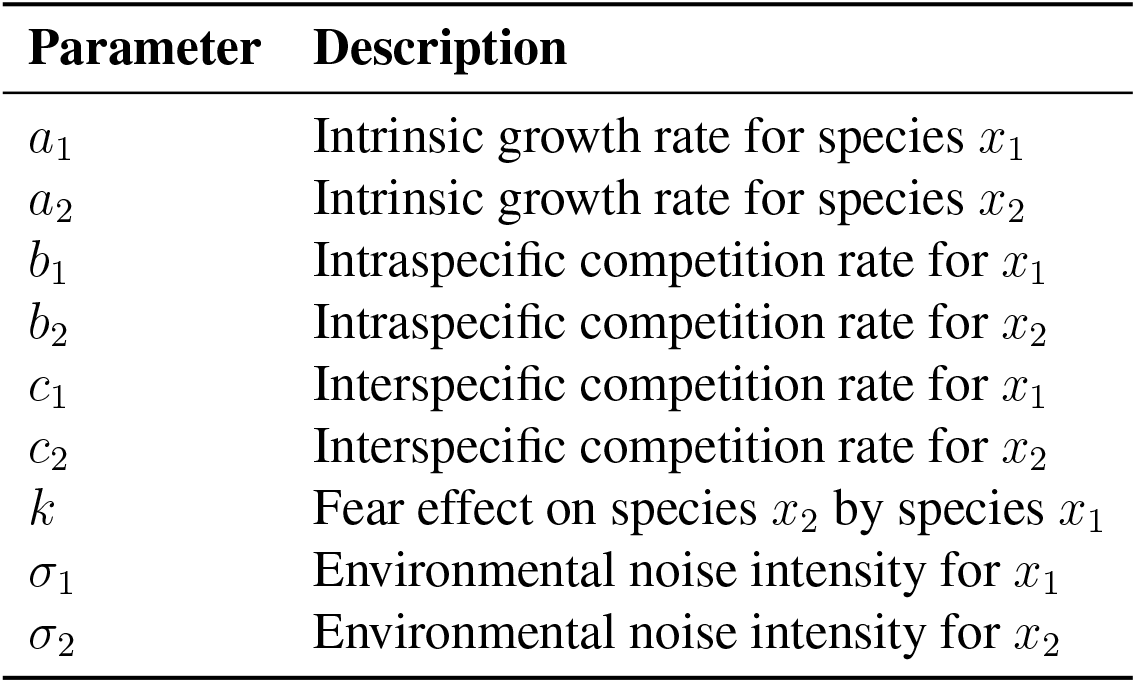
Model parameters and their biological descriptions for the stochastic fear-mediated two-species competition model.

## Results

### (a) Numerical validation of analytical thresholds and ecological case study

To complement the theoretical findings, we next perform a rigorous numerical exploration to analyze how environmental stochasticity reshapes the population dynamics of the system (3). We first use numerical simulations to validate the analytical extinction and persistence criteria established in Theorem 4 and Theorem 5, respectively (Figs. 1 and 2). We then run simulations for the case study of the Northern Spotted Owl and the Barred Owl using the calibrated parameters from [41]. We set *σ*_1_ = 0.2 and *σ*_2_ = 0.5. These values represent a setting in which the barred owl experiences weaker environmental variability, reflecting its broader ecological adaptability, whereas the spotted owl experiences stronger stochastic fluctuations associated with habitat degradation and reduced environmental tolerance [17, 42].

**Figure 1:**
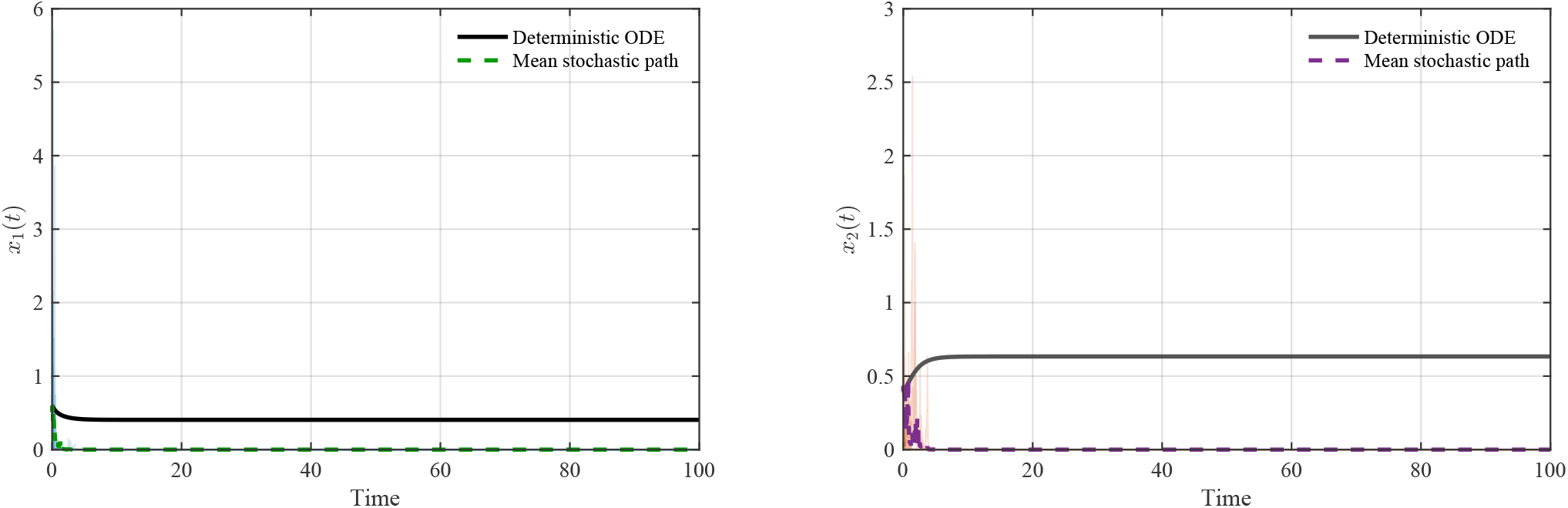
(Large Noise Drives Species to Extinction) Comparison of Deterministic (solid) vs mean stochastic (dashed) for *x*_1_ and *x*_2_ for *σ*_1_ = *σ*_2_ = 3.The other parameters used for the simulation are as follows: *a*_1_ = 1.0, *b*_1_ = 2.0, *c*_1_ = 0.3, *a*_2_ = 2.0, *b*_2_ = 2.0, *c*_2_ = 1.8, *k* = 0.005, and initial conditions *x*_1_(0) = 0.6 and *x*_2_(0) = 0.4.

**Figure 2:**
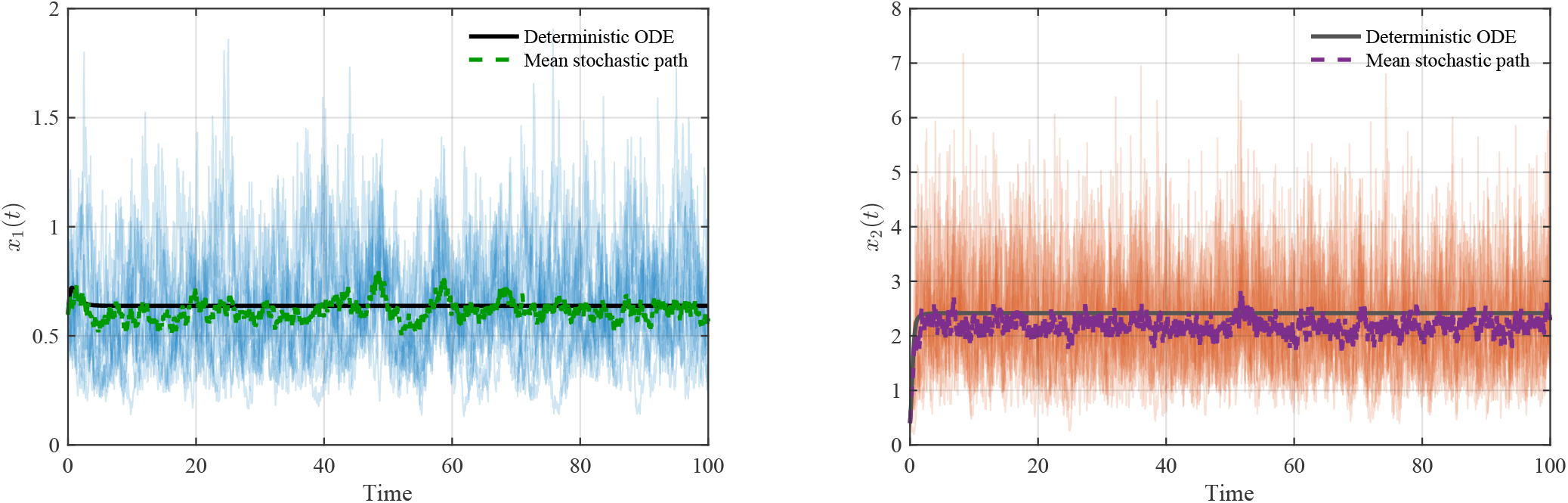
(Small Noise Maintains Species Co-existence) Comparison of Deterministic (solid) vs mean stochastic (dashed) for *x*_1_ and *x*_2_ for *σ*_1_ = 0.5 and *σ*_2_ = 1. The other parameters used for the simulation are as follows: *a*_1_ = 2.0, *b*_1_ = 2.0, *c*_1_ = 0.3, *a*_2_ = 6.0, *b*_2_ = 2.0, *c*_2_ = 1.8, *k* = 0.005, and initial conditions *x*_1_(0) = 0.6 and *x*_2_(0) = 0.4.

We observe that even under small environmental fluctuations, the spotted owl’s population *x*_2_ drives toward extinction earlier than predicted by the corresponding deterministic model [41] (Fig. 3). Stochasticity does not merely perturb the deterministic trajectory, but it can also heighten the extinction risk for ecologically vulnerable competitors.

**Figure 3:**
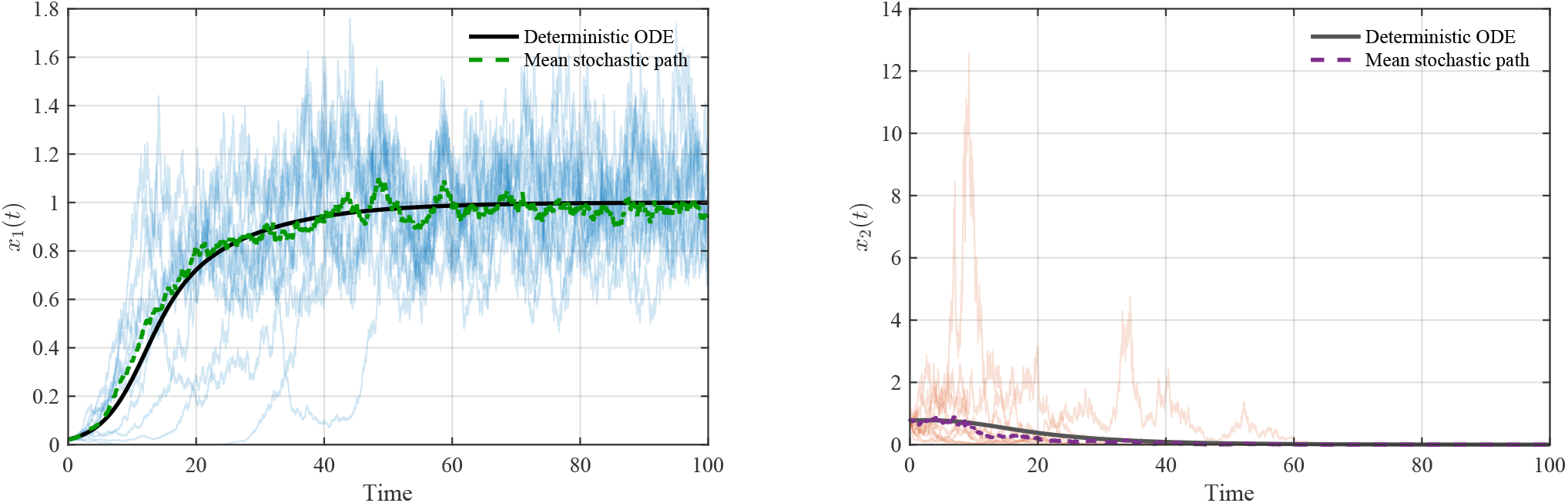
(Northern Spotted Owl and Barred Owl dynamics) Comparison of Deterministic (solid) vs mean stochastic (dashed) for Barred Owl (*x*_1_) and Northern Spotted Owl (*x*_2_) for *σ*_1_ = 0.2 and *σ*_2_ = 0.5. The other parameters used for the simulation are as follows: *a*_1_ = 0.547, *b*_1_ = 0.547, *c*_1_ = 0.307, *a*_2_ = 0.064, *b*_2_ = 0.064, *c*_2_ = 0.085, *k* = 4.725, and initial conditions *x*_1_(0) = 0.021 and *x*_2_(0) = 0.78 [41].

### (b) Environmental stochasticity can reverse deterministic competitive outcomes

Our simulations also reveal a stochastic reversal of competitive exclusion-type dynamics (Figs. 4 and 5). For the same parameter regime, the deterministic model (2) predicts that species *x*_1_ competitively excludes species *x*_2_. In contrast, under asymmetric environmental stochasticity with *σ*_1_ = 2.5 (stronger effect on *x*_1_) and *σ*_2_ = 0.1 (weaker effect on *x*_2_), the stochastic model (3) produces the opposite outcome: species *x*_2_ persists and ultimately excludes species *x*_1_ (Fig. 4). Similarly, in a second parameter regime, the deterministic model predicts that species *x*_2_ competitively excludes species *x*_1_, whereas asymmetric stochasticity with *σ*_1_ = 0.1 (weaker effect on *x*_1_) and *σ*_2_ = 2 (stronger effect on *x*_2_) reverses this outcome and allows species *x*_1_ to exclude species *x*_2_ (Fig. 5).

**Figure 4:**
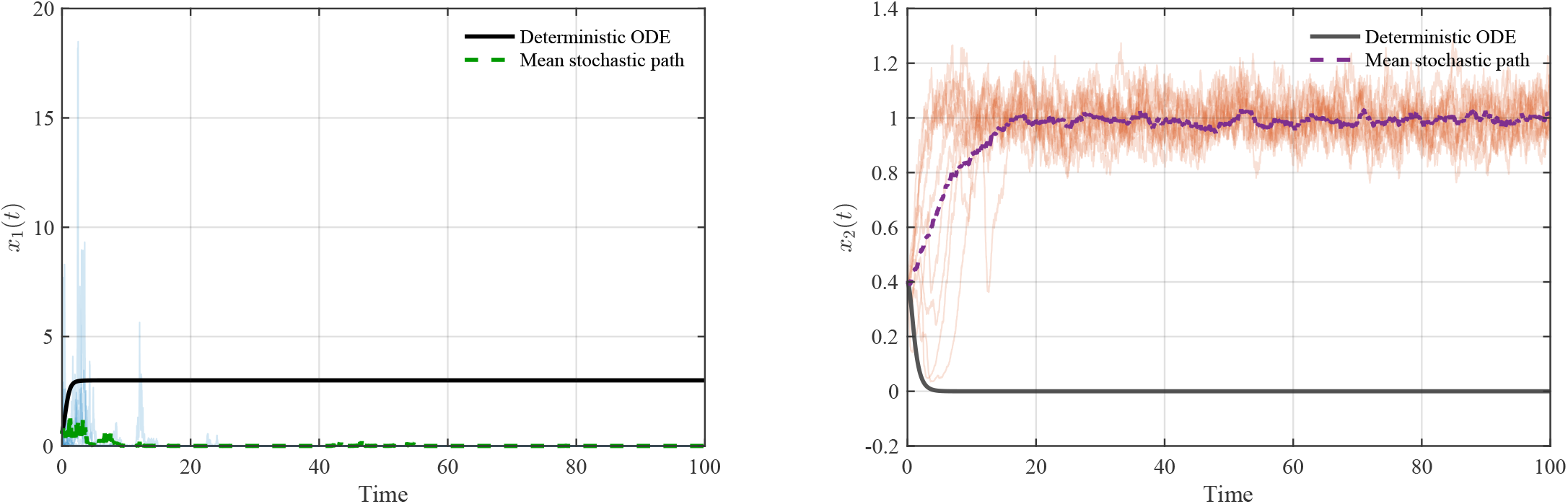
(CE reversal) Comparison of Deterministic (solid) vs mean stochastic (dashed) for *x*_1_ and *x*_2_ for *σ*_1_ = 2.5 and *σ*_2_ = 0.1.The other parameters used for the simulation are as follows: *a*_1_ = 3.0, *b*_1_ = 1.0, *c*_1_ = 0.5, *a*_2_ = 1.0, *b*_2_ = 1.0, *c*_2_ = 0.5, *k* = 1, and initial conditions *x*_1_(0) = 0.6 and *x*_2_(0) = 0.4.

**Figure 5:**
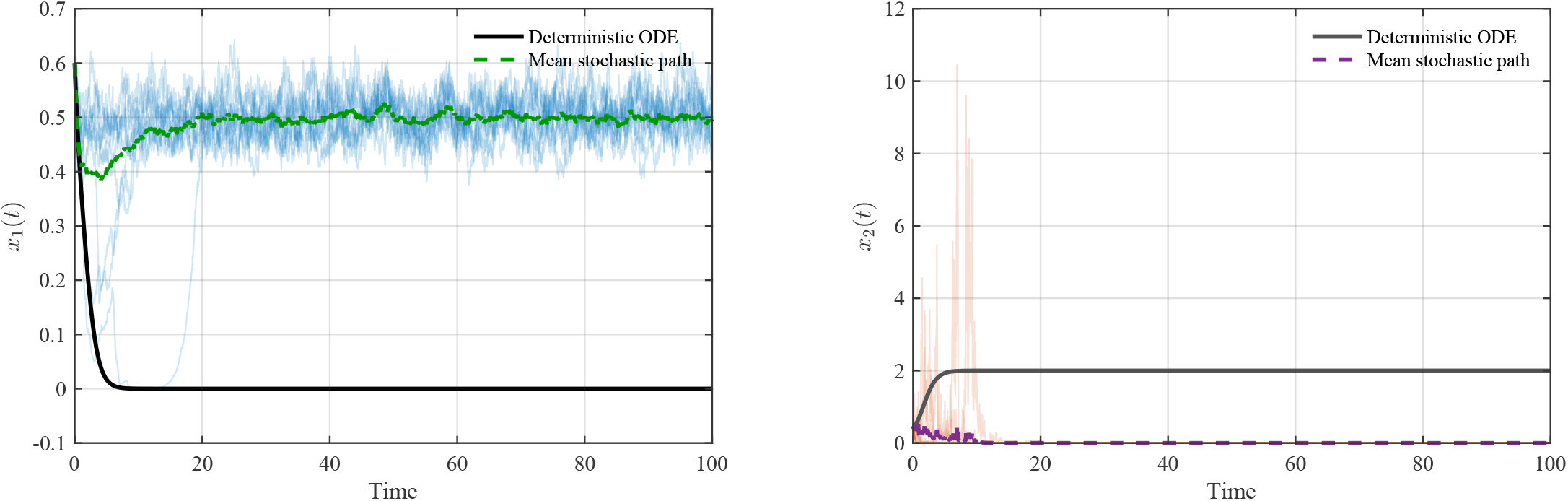
(CE reversal) Comparison of Deterministic (solid) vs mean stochastic (dashed) for *x*_1_ and *x*_2_ for *σ*_1_ = 0.1 and *σ*_2_ = 2.The other parameters used for the simulation are as follows: *a*_1_ = 1.0, *b*_1_ = 2.0, *c*_1_ = 1, *a*_2_ = 2.0, *b*_2_ = 1.0, *c*_2_ = 1.0, *k* = 0.5, and initial conditions *x*_1_(0) = 0.6 and *x*_2_(0) = 0.4.

This result is ecologically notable because environmental variability can reverse the deterministic outcome of competition. Stronger fluctuations can therefore drive the more noise-sensitive species to extinction, even when deterministic dynamics predict its competitive dominance.

### (c) Noise intensity reshapes coexistence dynamics

We next explore how different noise-intensity regimes alter the coexistence dynamics predicted by the deterministic system (Fig. 6). When species *x*_1_ experiences weak environmental stochasticity (*σ*_1_ = 0.1) and species *x*_2_ experiences strong environmental stochasticity (*σ*_2_ = 2), the dynamics of the stochastic system deviate substantially from deterministic coexistence (Fig. 6A). In this case, species *x*_1_ competitively excludes species *x*_2_ (Fig. 6A). In contrast, when the noise intensity acting on species *x*_2_ is reduced to a moderate level, *σ*_2_ = 1, species *x*_2_ persists, although its mean population density remains lower than that predicted by the deterministic model (Fig. 6B). In this regime, stochasticity indirectly benefits species *x*_1_ by suppressing the long-term abundance of its competitor (Fig. 6B).

**Figure 6:**
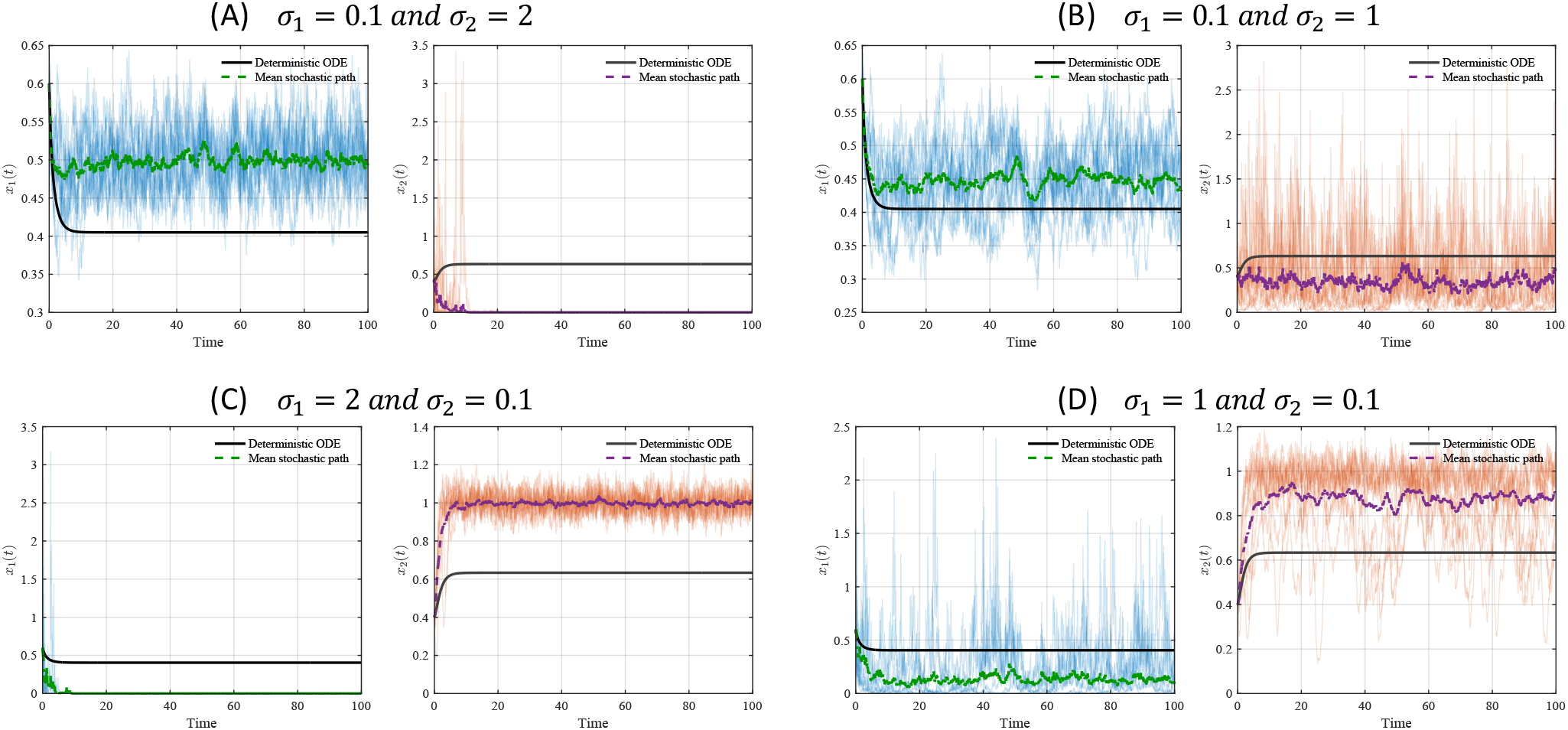
Comparison between deterministic and stochastic coexistence dynamics. Deterministic trajectories are shown by solid lines, and ensemble-mean stochastic trajectories are shown by dashed lines for *x*_1_ and *x*_2_ under different values of *σ*_1_ and *σ*_2_. The remaining parameters are *a*_1_ = 1.0, *b*_1_ = 2.0, *c*_1_ = 0.3, *a*_2_ = 2.0, *b*_2_ = 2.0, *c*_2_ = 1.8, *k* = 0.005, with initial conditions *x*_1_(0) = 0.6 and *x*_2_(0) = 0.4.

A symmetric pattern also emerges when species *x*_1_ experiences stronger environmental fluctuations. For the case of high noise intensity in species *x*_1_ (*σ*_1_ = 2) and weak noise intensity in species *x*_2_ (*σ*_2_ = 0.1), the stochastic system (3) produces competitive exclusion of species *x*_1_ by species *x*_2_ (Fig. 6C). However, when the noise intensity acting on species *x*_1_ is reduced to a moderate level, *σ*_1_ = 1, species *x*_1_ persists but at a reduced mean density relative to the deterministic coexistence state (Fig. 6D). In this scenario, stochasticity indirectly benefits species *x*_2_ by reducing the competitive pressure from species *x*_1_ (Fig. 6D). Together, these simulations show that deterministic coexistence may be robust to moderate stochasticity, but sufficiently strong species-specific fluctuations can transform coexistence into competitive exclusion.

### (d) Species-specific noise thresholds structure persistence and extinction

To examine the interplay between fear and stochasticity in greater detail, we then vary both environmental stochasticity *σ*_*i*_ (for *i* = 1, 2) and the fear parameter, *k*, and output the long-time population means ⟨*x*_1_⟩ and ⟨*x*_2_⟩ (Figs. 7A–B and 8A–B). For better visualization, we also output a heatmap that classifies the extinction (red) and persistence (blue) dynamics for each species (Figs. 7C–D and 8C–D).

**Figure 7:**
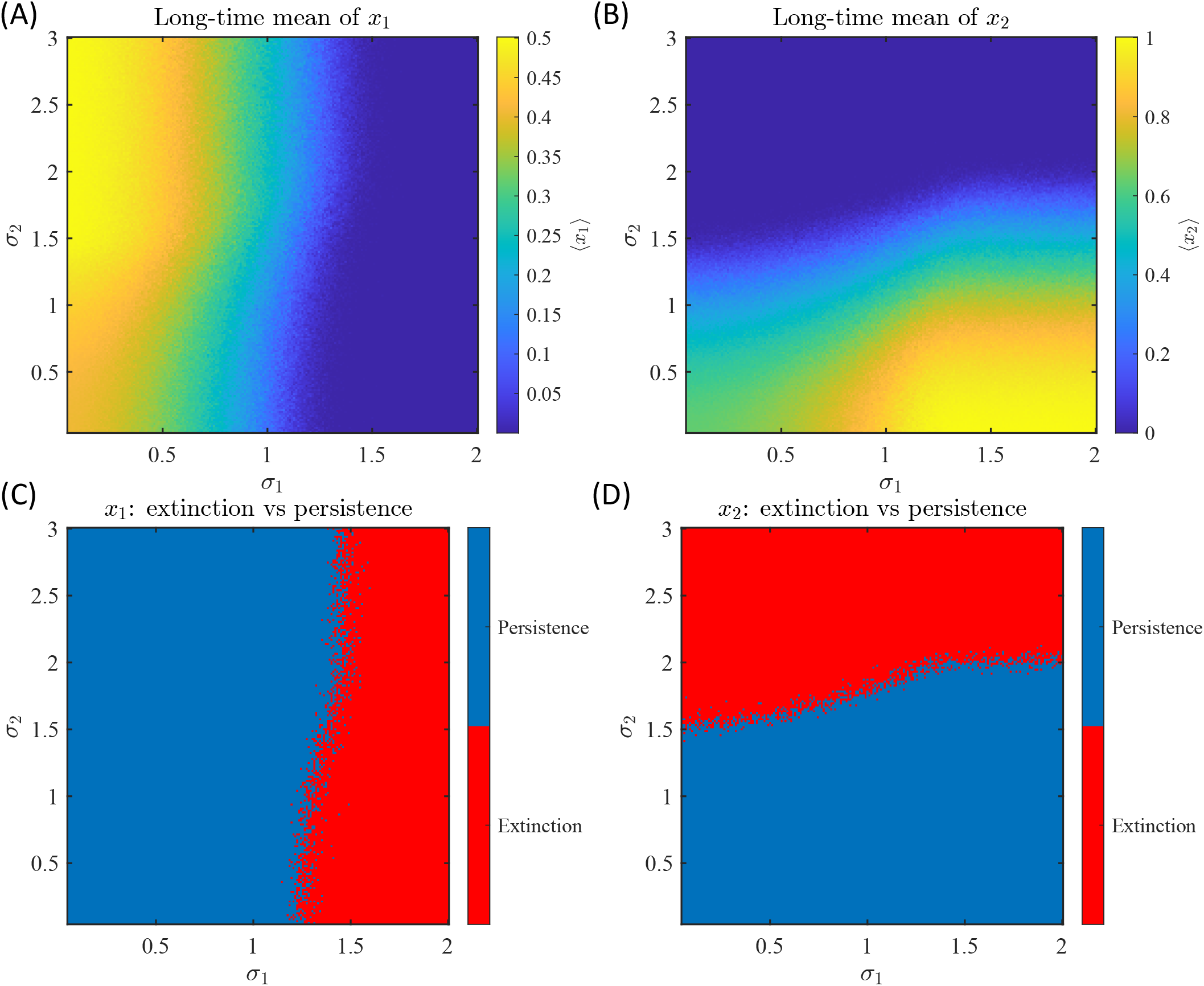
Effects of species-specific environmental stochasticity on long-time population means and persistence outcomes. Heatmaps in the (*σ*_1_, *σ*_2_)-plane show the long-time mean densities of (A) ⟨*x*_1_⟩ and (B) ⟨*x*_2_⟩, together with the corresponding extinction–persistence classifications for (C) *x*_1_ and (D) *x*_2_. The remaining parameters are *a*_1_ = 1.0, *b*_1_ = 2.0, *c*_1_ = 0.3, *a*_2_ = 2.0, *b*_2_ = 2.0, *c*_2_ = 1.8, *k* = 0.005, with initial conditions *x*_1_(0) = 0.6 and *x*_2_(0) = 0.4.

In the (*σ*_1_, *σ*_2_) plane, the long-time mean of species *x*_1_ decreases primarily with increasing *σ*_1_. This indicates that the persistence of species *x*_1_ is strongly controlled by its own environmental noise intensity (Fig. 7A). For small to moderate values of *σ*_1_, species *x*_1_ maintains a positive long-time mean over a broad range of *σ*_2_. In contrast, sufficiently large *σ*_1_ values, approximately greater than 1.5, drive the system into an extinction regime for *x*_1_. A similar pattern is observed in the binary extinction-persistence heatmap. The transition boundary for *x*_1_ is almost vertical, suggesting that *σ*_1_ is the dominant stochastic driver of extinction for this species (Fig. 7C).

In contrast, the long-time mean of *x*_2_ is more strongly reduced by increasing *σ*_2_, with persistence occurring mostly below a critical *σ*_2_ threshold (Fig. 7B). The extinction–persistence boundary for *x*_2_ is therefore primarily organized along the *σ*_2_ direction (Fig. 7D). However, this boundary also exhibits a slight dependence on *σ*_1_, suggesting that noise acting on the competing species can indirectly influence the persistence of *x*_2_ by altering competitive pressure. Overall, these simulations show that each species is most vulnerable to stochastic perturbations acting directly on its own growth dynamics. Moreover, cross-species stochastic effects remain important because competition couples the long-term outcomes of the two populations.

### (e) Fear and stochasticity jointly constrain species persistence

The second set of heatmaps explores the joint effect of the fear parameter *k* and a common noise intensity *σ*, where *σ*_1_ = *σ*_2_ = *σ*. The long-time mean of *x*_1_ is mostly independent of increasing *k* but declines sharply as *σ* increases (Fig. 8A). A similar trend can also be observed in the extinction– persistence heatmap for *x*_1_ (Fig. 8C). Fig. 8C confirms this interpretation, as the boundary between persistence and extinction is almost horizontal in the (*k, σ*) plane, showing that stochastic intensity, rather than fear strength, determines whether *x*_1_ persists.

**Figure 8:**
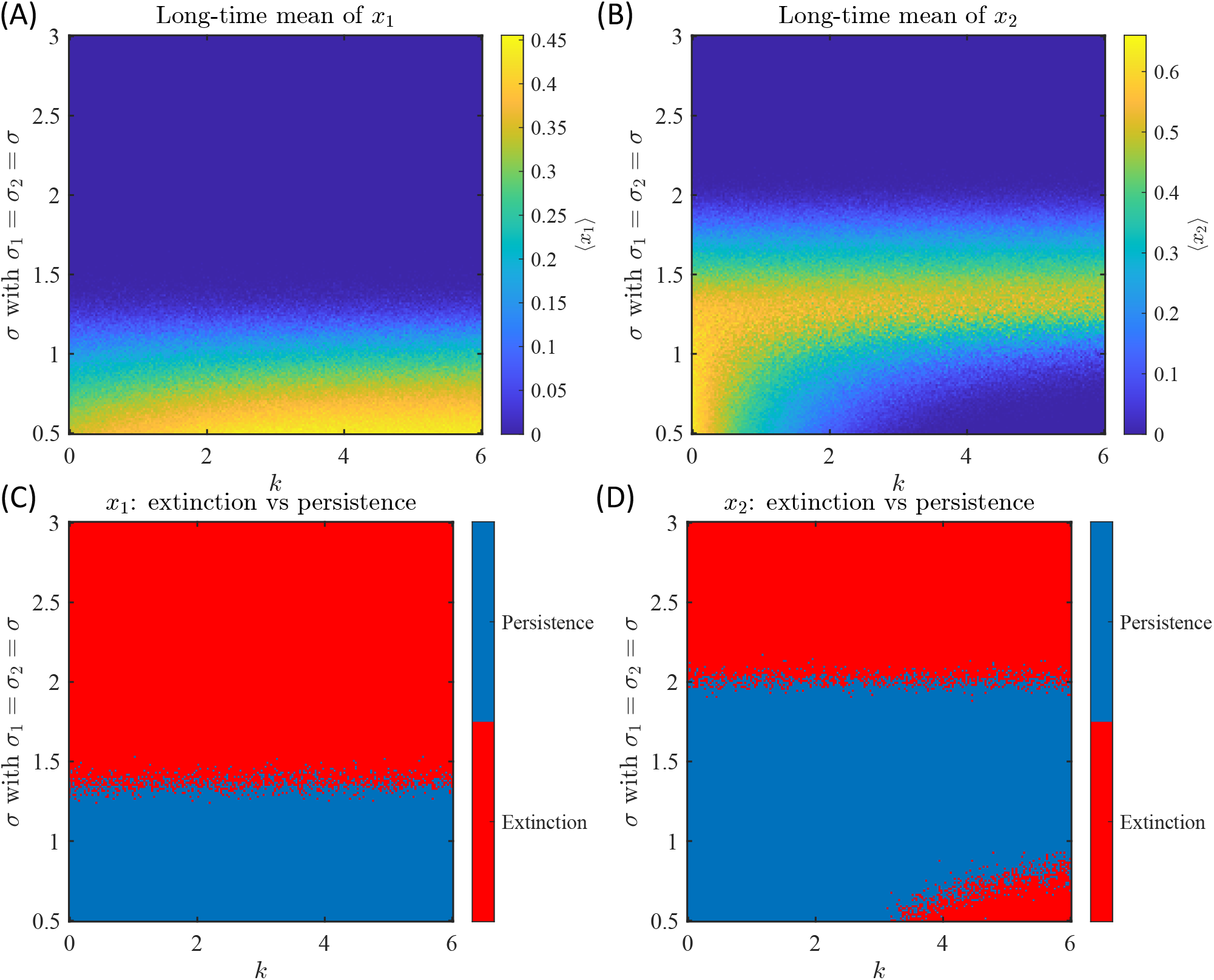
Joint effects of fear strength and environmental stochasticity on long-time population means and persistence outcomes. Heatmaps in the (*k, σ*)-plane, with *σ*_1_ = *σ*_2_ = *σ*, show the long-time mean densities of (A) ⟨*x*_1_⟩ and (B) ⟨*x*_2_⟩, together with the corresponding extinction–persistence classifications for (C) *x*_1_ and (D) *x*_2_. The remaining parameters are *a*_1_ = 1.0, *b*_1_ = 2.0, *c*_1_ = 0.3, *a*_2_ = 2.0, *b*_2_ = 2.0, *c*_2_ = 1.8, with initial conditions *x*_1_(0) = 0.6 and *x*_2_(0) = 0.4.

For *x*_2_, however, the interaction between fear and noise is more pronounced. The long-time mean of *x*_2_ is highest when both *k* and *σ* are small to moderate, but declines as either the fear parameter or noise intensity increases (Fig. 8B). At low environmental noise, *x*_2_ can persist across a broad range of *k*, although increasing fear gradually reduces its long-time mean. At higher values of *σ*, the persistence region narrows substantially, and large values of *k* push *x*_2_ toward extinction even when the noise level is not sufficient by itself to eliminate the species. The extinction–persistence heatmap for *x*_2_ therefore reveals a coupled threshold structure: environmental noise lowers the species’ tolerance to fear, while stronger fear reduces the amount of stochastic variability the species can withstand (Fig. 8). Together, these heatmap results support the analytical prediction that extinction and persistence are governed by threshold-like conditions involving both environmental noise and fear intensity.

## Conclusion

In this work, we develop a novel stochastic differential equation framework (3) that captures the interplay between competitive interactions of two species, the non-consumptive fear effect, and environmental stochasticity. From an ecological perspective, the parameter *σ*_*i*_ in the (3) represents the intensity of environmental variability experienced by species *x*_*i*_. Such variability may arise from random fluctuations in temperature, precipitation, food availability, habitat quality, disturbance frequency, or other environmental drivers that alter birth, death, and interaction rates. There are several ecological case studies that highlight the significance of stochasticity and the effect of fear on population dynamics (stochasticity effect: [26–31]; Fear effect: [17–21]). Additionally, empirical studies suggest that environmental stochasticity (in the form of how changes in the habitat and climate) and fear effects can affect the population dynamics of competing species [18, 21, 33, 34]. In this work, we incorporate multiplicative environmental noise into the current competing-species model to capture realistic ecological uncertainties [35–40]. This approach enhances our understanding of how random fluctuations and non-consumptive effects shape long-term population dynamics. The stochastic formulation allows us to explore how random environmental disturbances modify deterministic predictions, identify thresholds for persistence and extinction, and gain insight into the resilience of ecological systems that are subject to unpredictable changes.

We first establish the well-posedness of the system (3) in 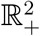 almost surely (Lemma 1, 2 and Theorem. 3). We use Lyapunov-type estimates to control the solution’s growth and prove almost sure eventual boundedness (Theorem 3). This boundedness result shows that the solution trajectories model remains mathematically well-defined and biologically admissible for all future time. We then derive sharp threshold conditions to characterize the system’s long-time dynamics and how environmental stochasticity reshapes extinction and persistence. In particular, Theorem 4 provides a sufficient condition under which environmental stochasticity dominates intrinsic growth, leading to almost sure exponential extinction of both species (Fig. 1). This highlights the destabilizing role of sufficiently strong noise, even in parameter regimes where the corresponding deterministic system admits a locally stable coexistence equilibrium. Essentially, the environmental stochasticity lowers the effective growth rates, and when this reduction dominates the intrinsic growth contributions, both species decay exponentially almost surely.

Conversely, under appropriate boundedness assumptions on the dominant competitor, we prove a persistence criterion for the fearful species in Theorem 5 (Fig. 2). The derived threshold *λ*_2_ explicitly quantifies the balance between fear-induced fitness suppression, interspecific competition, and stochastic fluctuations. Notably, this condition reveals how the fear parameter *k* and the noise intensity *σ*_2_ jointly regulate long-term survival. Together, these results show that environmental stochasticity does not simply perturb the deterministic dynamics; it can qualitatively alter the outcome by shifting the system between stochastic extinction and mean persistence.

Our theoretical results motivate a detailed study of the effects of environmental stochasticity. We perform a rigorous numerical analysis to validate our theoretical results (Figs. 1 2, and 6) as well as uncover key ecological dynamics such as competition exclusion reversal (Figs. 5 and 4). Our heatmap analysis shows that increasing environmental stochasticity can shift a population from persistence to extinction even when deterministic dynamics would otherwise allow long-term survival (Figs. 7 and 8).

The distinct roles of *σ*_1_ and *σ*_2_ suggest that species-specific vulnerability to environmental variation is crucial [17, 42, 43] (Fig. 7). Species *x*_1_ persists when *σ*_1_ remains below a critical range, largely independent of the stochasticity affecting *x*_2_. Biologically, this implies that a competitively stronger or less fear-affected species may remain viable unless environmental fluctuations directly impair its own demographic rates. In contrast, species *x*_2_ is more vulnerable because its growth is already reduced by the fear-mediated effect of *x*_1_. Therefore, stochasticity acting on *x*_2_ does not operate in isolation; rather, it compounds an existing behavioral or physiological cost. This produces a lower effective growth margin, making *x*_2_ more susceptible to extinction under environmental variability. The (*k, σ*) heatmaps analysis reveals that fear alone can reduce the long-time mean of *x*_2_, but its most severe effects occur when combined with environmental stochasticity (Fig. 8). This finding is ecologically significant, suggesting that a fearful competing species may still persist under relatively stable environmental conditions, but the same species may be pushed toward extinction when random environmental fluctuations further reduce its growth opportunities.

These results highlight a biologically relevant mechanism by which non-consumptive effects and environmental variability jointly reshape coexistence outcomes. In deterministic settings, fear may reduce population density without necessarily causing extinction. However, under stochastic environmental conditions, fear can reduce the buffer between persistence and extinction. As a result, populations exposed to strong behavioral suppression may become less resilient to climatic variability and habitat disturbance. From a biological conservation manager’s perspective, these findings imply that species exposed to both competitive pressure and fear effects may require lower environmental variability, better habitat quality, or reduced disturbance to persist [36, 37, 44, 45]. Management strategies that only reduce direct mortality may be insufficient if non-consumptive effects continue to suppress growth or reproduction. Thus, the heatmap analysis provides a practical way to identify regions of parameter space where persistence is robust, where extinction is likely, and where small increases in fear or environmental variability can trigger a qualitative transition in long-time population dynamics.

Our analysis also has implications that extend beyond the model’s specific mathematical structure. A key takeaway is that environmental stochasticity can impact coexistence outcomes, with this effect being highly dependent on the ecological context in which species interact. Rather than serving as a uniform destabilizing force, stochasticity interacts with competition and fear-induced suppression to alter the effective boundary between persistence and extinction. This finding is particularly relevant for empirical studies, as many natural populations experience a combination of behavioral stressors and environmental variability. Such variability includes fluctuating resources, temperature extremes, habitat disturbances, and predator-induced changes in activity or foraging behavior. Consequently, the results also suggest that extinction risk cannot be assessed solely on the basis of average interaction strengths; instead, it may be influenced by how variability enhances or diminishes crucial biological mechanisms. In this regard, the threshold established in this study serves as a conceptual bridge between mathematical persistence theory and ecological risk assessment, demonstrating how environmental noise can transform a system that is expected to persist deterministically into one that exhibits extinction-like dynamics.

Altogether, our results provide a rigorous description demonstrating that environmental stochasticity can either suppress populations to extinction or allow them to persist, depending on its interaction with behavioral effects. All of these findings offer a mechanistic and mathematically explicit framework for understanding how non-consumptive interactions and randomness shape ecological outcomes. Thereby, the insights from this work have potential implications for the management of competing species under uncertain environmental conditions.

## Material and Methods

### (a) Mathematical framework and analytical results

In this work, we aim to study the long-term dynamics of the stochastic fear-mediated competition model (3). The model (3) extends the deterministic fear-mediated competition system (2) by incorporating environmental stochasticity through multiplicative noise terms. In this formulation, *x*_1_(*t*) and *x*_2_(*t*) denote the population densities of the two competing species, while the parameters *a*_*i*_, *b*_*i*_, and *c*_*i*_ describe intrinsic growth, intraspecific competition, and interspecific competition, respectively. The parameter *k* measures the strength of the fear effect on species *x*_2_, and *σ*_*i*_(*t*) denotes the intensity of environmental stochasticity acting on species *x*_*i*_.

The deterministic system (2) provides the baseline ecological dynamics in the absence of environmental fluctuations, whereas the stochastic model (3) is used to examine how random environmental perturbations modify these deterministic predictions. This comparison allows us to determine whether stochasticity preserves coexistence, accelerates extinction, or reverses deterministic competitive outcomes. Throughout the analysis, all initial conditions are assumed to be positive, since the state variables represent population densities.

In this work, the analytical results focus on three mathematical properties of the stochastic system: well-posedness, boundedness, and long-term extinction or persistence. All of the Lemma and Theorem proof details are given in Appendix A. In the main text, we will state the key results:

#### Lemma 1.

*Let us assume* (*x*_1_(*t*), *x*_2_(*t*)) *are the solution to the system* (3), *then x*_1_ *and x*_2_ *is eventually bounded a*.*s*.

#### Lemma 2.

*For the system* (3), *the local solution* (*x*_1_(*t*), *x*_2_(*t*)) *exists a*.*s. for t* ∈ [0, *τ*_*e*_) *for positive initial data: x*_1_(0) *>* 0, *x*_2_(0) *>* 0, *where τ*_*e*_ *is defined as the explosion time*.

#### Theorem 3.

*For the system* (3) *with positive initial conditions, there exits a unique solution* (*x*_1_(*t*), *x*_2_(*t*)). *Moreover, this solution will remain in* 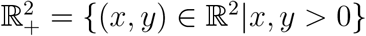 *with probability 1*.

These results establish that the stochastic model is mathematically well-defined and biologically meaningful. This means that positive initial population densities remain positive, and the long-term trajectories remain bounded almost surely.

#### Theorem 4.

*The interior equilibrium* 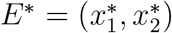 *of the deterministic system* (2) *is locally asymptotically stable if* 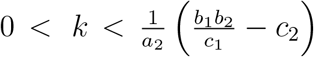. *Further assume that the noise intensity satisfies* 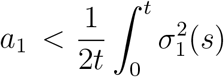 d*s and* 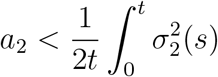 d*s. Then the solutions x*_1_(*t*) *and x*_2_(*t*) *of the stochastic system* (3) *go to extinction exponentially almost surely*.

The extinction theorem identifies stochastic regimes in which environmental fluctuations exceed intrinsic population growth and drive both species to extinction. This provides a direct mathematical explanation for cases in which stochastic dynamics exert stronger extinction pressure than the deterministic model predicts.

#### Theorem 5.

*Assume that the positive solution of* (3) *satisfies* 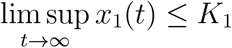 *a.s for some K*_1_ *>* 0.

*Define*

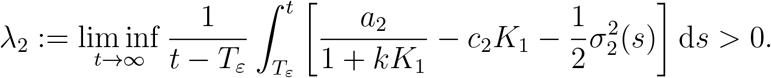

*Then*,

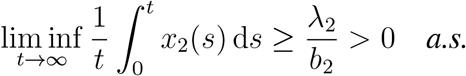

The persistence theorem provides a long-time-average criterion for the persistence of species *x*_2_. Ecologically, this condition shows that species *x*_2_ persists when its effective stochastic growth rate remains positive after accounting for fear, competition from species *x*_1_, and environmental noise. Thus, the theorem provides an explicit threshold-type connection between fear intensity and stochastic forcing.

Together, these analytical results provide the mathematical foundation for the numerical simulations. The extinction theorem is used to identify noise regimes in which stochasticity can drive species to extinction, while the persistence theorem motivates the numerical exploration of coupled fear-noise thresholds.

### (b) Numerical simulations

We performed the numerical simulation in `MATLAB` for both the deterministic and stochastic versions of system (3) to validate the theoretical results [46]. To solve the ODE, we used the built-in `MATLAB` solver `ode45`, which is based on an adaptive Runge–Kutta (4,5) method. We set the relative and absolute tolerances to `RelTol`= 10^−9^ and `AbsTol` = 10^−10^, respectively to ensure high numerical precision. We use the deterministic ODE system (2) to serve as a reference for comparison with the stochastic system. For the stochastic model (3), simulations were performed using the Euler-Maruyama (EM) method [47, 48]. The EM method is the standard numerical approximation scheme for stochastic differential equations with multiplicative environmental noise. For the simulation, we used a fixed time step of Δ*t* = 0.01 over the simulation interval to accurately capture stochastic fluctuations while maintaining computational efficiency. The Brownian motion terms were approximated using independent normally distributed random variables generated with `MATLAB`’s `randn` function. These random variables are then scaled appropriately by the square root of the time step. Since the model describes population densities, the positivity of solutions was numerically preserved by setting any negative values arising from discretization errors to zero after each iteration.

To examine the impact of environmental stochasticity, we conducted multiple stochastic realizations using Monte Carlo simulations [49]. We generated several independent sample paths to visualize random fluctuations around the deterministic dynamics, while the average stochastic trajectory was computed as the mean across all realizations. This method enables a direct comparison among deterministic solutions, individual stochastic sample paths, and ensemble-mean behavior. All figures were created using `MATLAB`’s high-resolution plotting tools and were exported in vector format for publication-quality presentation.

### (c) Parameter sweeps and persistence-extinction classification

To examine how environmental stochasticity reshapes long-term population outcomes, we performed systematic parameter sweeps over the noise intensities *σ*_1_ and *σ*_2_. For each pair (*σ*_1_, *σ*_2_), the stochastic system (3) was simulated over a long time interval, and the corresponding long-time mean densities ⟨*x*_1_⟩ and ⟨*x*_2_⟩ were computed. These values were then used to generate heatmaps showing how each species responds to direct and indirect stochastic forcing.

We also explored the joint effect of fear and environmental variability by varying the fear parameter *k* together with a common noise intensity *σ* (by setting *σ*_1_ = *σ*_2_ = *σ*). This analysis was done to identify regimes in which environmental noise amplifies the population-level effects of fear. In particular, these simulations test the analytical prediction that persistence of species *x*_2_ depends on a coupled threshold involving both the fear parameter and the stochastic noise intensity.

We also explored the binary classification of persistence and extinction. A species was classified as persistent if its long-time mean density remained above a small numerical threshold and was classified as extinct if its long-time mean density fell below that threshold. This classification was used only as a numerical summary of the long-time simulations; the analytical extinction and persistence results remain stated in the almost-sure and long-time average senses.

### (d) Ecological case study

We also simulated the model (3) using calibrated parameters from the Northern Spotted Owl-Barred Owl competition case study [41]. For this ecological case study, the environmental stochasticity was incorporated to represent random fluctuations in ecological conditions, including variation in habitat quality, resource availability, and disturbance intensity. For this case study, we used *σ*_1_ = 0.2 and *σ*_2_ = 0.5, representing a scenario in which the barred owl is less sensitive to environmental fluctuations, while the northern spotted owl experiences stronger stochastic effects. This choice is consistent with ecological evidence that barred owls are more adaptable across heterogeneous habitats, whereas northern spotted owls are more strongly affected by habitat degradation and environmental change [17, 42]. The stochastic simulations were then compared with the deterministic model to evaluate whether environmental variability accelerates the decline of the northern spotted owl relative to deterministic predictions.

## Data availability

No original data were used in this manuscript, and all search terms and articles from the literature review are described and cited.

## Appendix A. Mathematical Model and Results

### Mathematical Model

ODE Model:

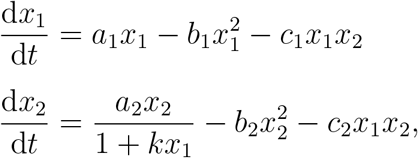

Stochastic Model

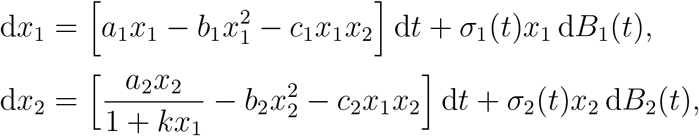

### Mathematical Results

#### Lemma 1.

*Proof*. Consider the following SDE,

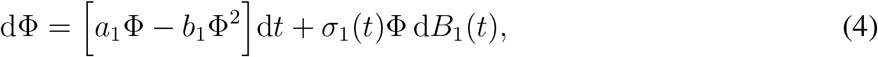

where Φ(*t*) denotes the solution. Note that the solution for (4) exists globally and the proof follows standard techniques [50]. Hence, *τ* = ∞. Moreover if

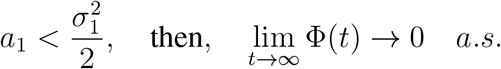

In the case when 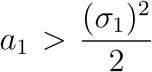, the persistence result is expressed in the form of a mean, which means

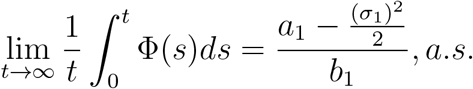

Similarly, we have 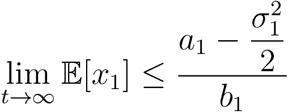, we obtain that the *x*_1_ is eventually bounded.

#### Lemma 2.

*Proof*. Define *x*_1_(*t*) = *e*^*u*(*t*)^ and *x*_2_(*t*) = *e*^*v*(*t*)^. Hence, *u*(*t*) = ln(*x*_1_(*t*)) and *v*(*t*) = ln(*x*_2_(*t*)). On using the Itô’s formula, we have the following system with initial conditions *u*(0) = ln *x*_1_(0) and *v*(0) = ln *x*_2_(0):

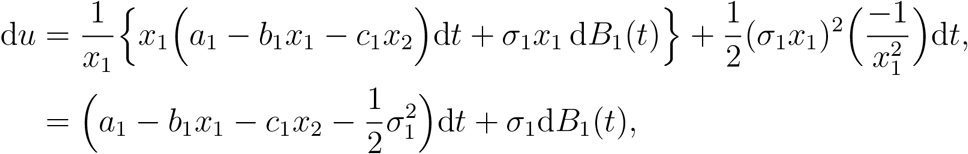

Similarly,

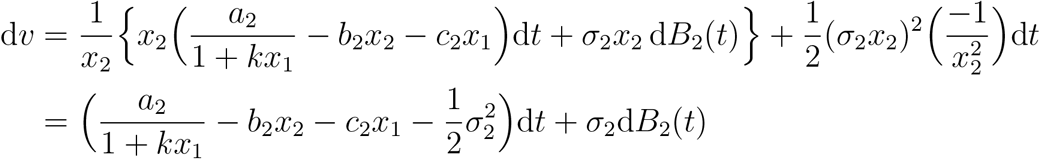

Given that *x*_1_(*t*) = *e*^*u*(*t*)^ and *x*_2_(*t*) = *e*^*v*(*t*)^, we have the following system with initial conditions: *u*(0) = ln(*x*_1_(0)) and *v*(0) = ln(*x*_2_(0)).

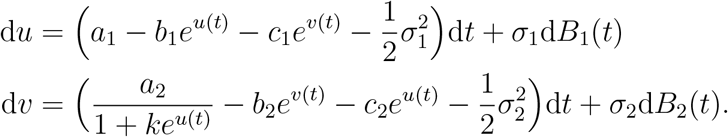

Since the coefficients of the above system satisfy the local Lipschitz conditions, for given positive initial conditions, we have a unique maximum local solution *u*(*t*) and *v*(*t*) on the interval [0, *τ*_*e*_). From the Itô’s formula [50], and the fact that *x*_1_(*t*) = *e*^*u*(*t*)^ and *x*_2_(*t*) = *e*^*v*(*t*)^, we have positive local solution for positive initial conditions.

#### Theorem 3.

*For the system* (3) *with positive initial conditions, there exits a unique solution* (*x*_1_(*t*), *x*_2_(*t*)). *Moreover, this solution will remain in* 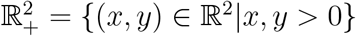 *with probability 1*.

*Proof*. Let

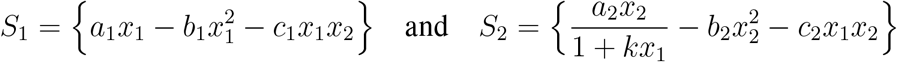

denotes the source terms for system (3).

Let 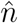 (a large number) such that the initial data lies inside 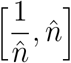, and also define the stopping time as follows:

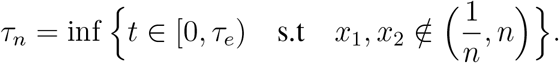

By definition, *τ*_*n*_ is increasing sequence and 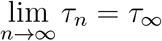 whenever *τ*_∞_ ≤ *τ*_*e*_ a.s.

To prove the theorem, it suffices to show that *τ*_*e*_ = ∞. For the proof, we will make use of the method of contradiction. Let’s assume that the above statement is false, then there exists a 0 *< T <* ∞ and *ϵ* (small number ∈ (0, 1)) such that ℙ{*τ*_∞_ ≤ *T*} *> ϵ*. This implies that 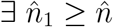 such that

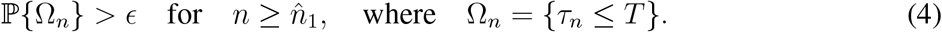

Define a *C*^2^ function 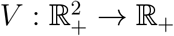 as follows:

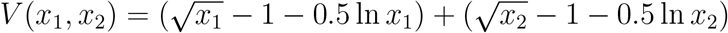

For 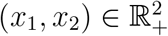 and applying the Itô formula [50, 51], we have

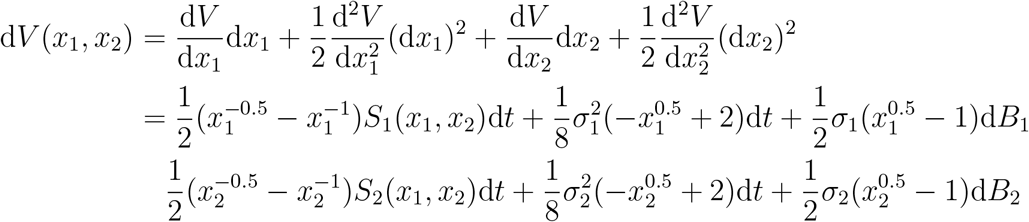

From, Lemma 1, we have the following inequality

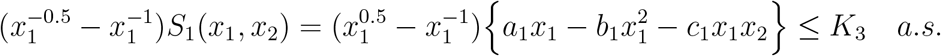

For this inequality, we assume that sufficient time has passed for the bound to be achieved. This time may depend on the initial condition; for larger initial conditions, more time may be needed for absorption. Similarly, we have

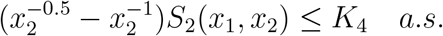

Hence, we have

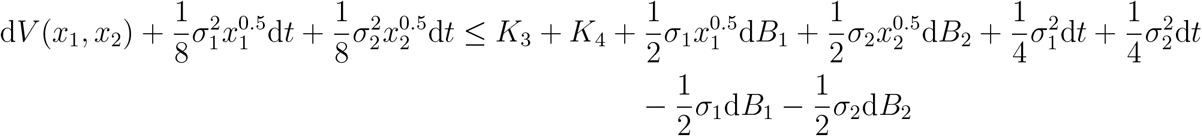

By applying positivity, assumption on 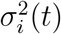 and integrating the equation from 0 to 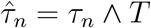, and then taking the expectation on both sides, we have,

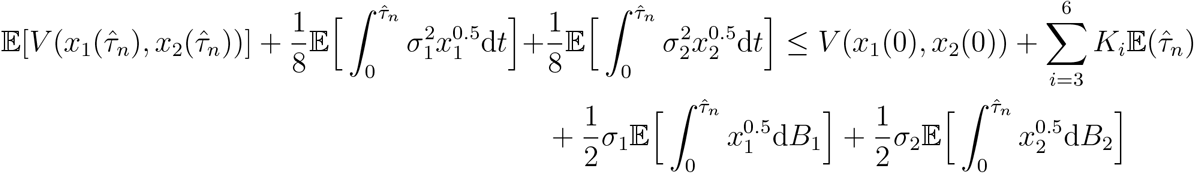

We can also find the bounds for the following term using the fact that this Itô integral is an embedding of ℒ^2^ ↪ ℒ^1^, Itô isometry, and application of Holder’s and Young’s inequality, we get

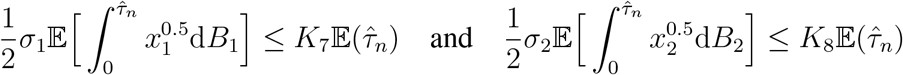

Hence, we have 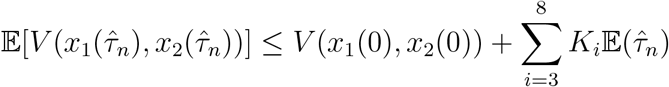.

Recall, since ℙ(Ω_*n*_) ≥ *ϵ*. Hence, from (4), we have for *w* ∈ Ω_*n*_, there exists some *i* such that *x*_*i*_(*τ*_*n*_, *w*) for *i*=1,2 and equals to *n* or 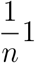. Hence, 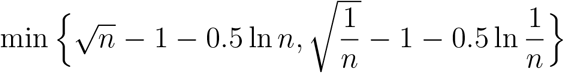 is lower bound for 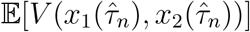. For indicator function 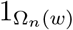, we have

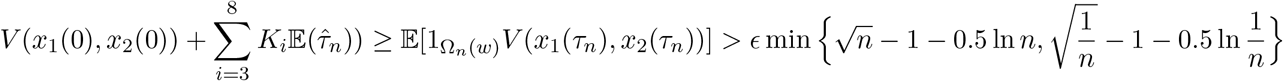

As *n* → ∞, we have a contradiction: 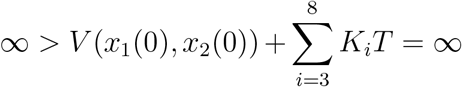, and this completes the proof.

#### Theorem 4.

*The interior equilibrium* 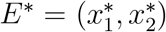 *of the deterministic system* (2) *is locally asymptotically stable if* 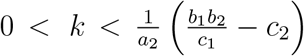. *Further assume that the noise intensity satisfies* 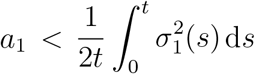 *and* 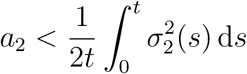. *Then the solutions x*_1_(*t*) *and x*_2_(*t*) *of the stochastic system* (3) *go to extinction exponentially almost surely*.

*Proof*. On evaluating the Jacobian matrix evaluated at *E*^∗^, we have 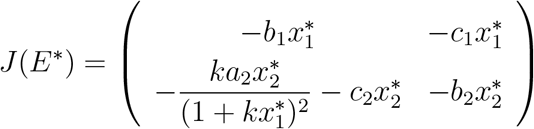.

To show that the equilibrium *E*^∗^ is local stable, it is enough to show that *tr*(*J*(*E*^∗^)) *<* 0 and det(*J*(*E*^∗^)) *>* 0. On computation, we have

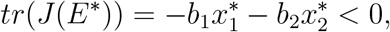

and

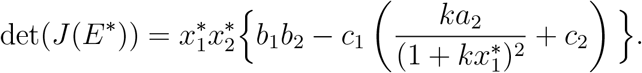

Therefore, on choosing *k* is such that,

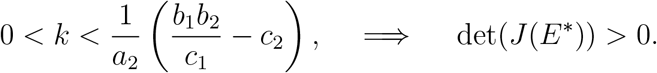

Hence, the interior equilibrium *E*^∗^ is locally asymptotically stable [23]. Now, applying the Itô’s formula for system (3) and using the comparison result [52], we have

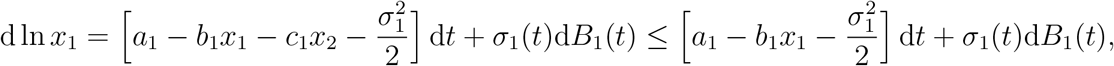

On integrating from 0 to *t*, we have

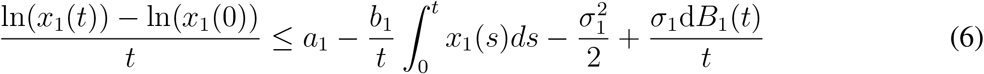

On using the strong law of large numbers, we have

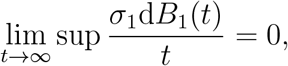

and

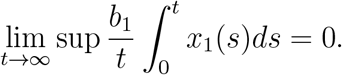

Hence, under the assumption 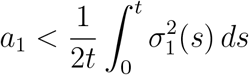, we have

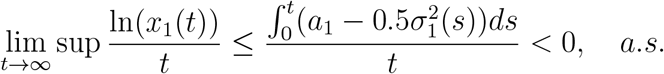

So, *x*_1_ goes to extinction, a.s. Similarly, on using the comparison theorem [52] and the fact that *x*_1_ goes to extinction, a.s, thus the other equation for *x*_2_ reduces to the following

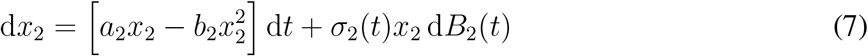

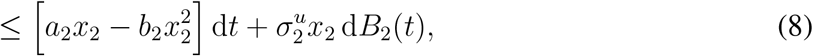

where 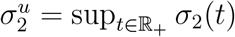. On using the Itô’s formula for the above equation, we have

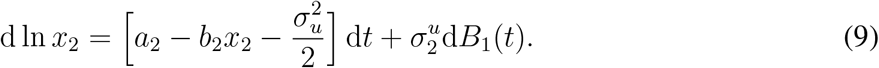

Similarly, under the assumption 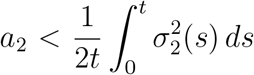 and from (7), we have 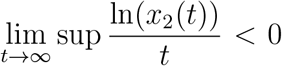 a.s. Thus, *x*_2_ goes to extinction a.s.

#### Theorem 5.

*Assume that the positive solution of* (3) *satisfies* 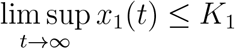 *a*.*s for some K*_1_ *>* 0. *Define*

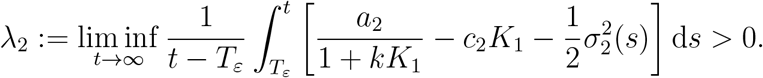

*Then*,

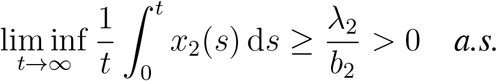

*Proof*. On using the assumption on *x*_1_(*t*), we have for every *ε >* 0 there exists *T*_*ε*_ such that *x*_1_(*t*) ≤ *K*_1_ + *ε*, when *t* ≥ *T*_*ε*_, a.s. Hence, on using the Itô’s formula on ln *x*_2_, for all sufficiently large *t*, we have

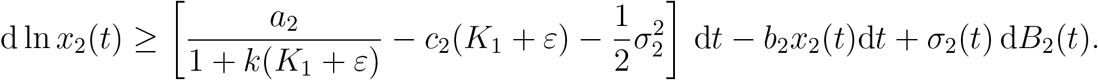

On integrating from *T*_*ε*_ to *t*, we have

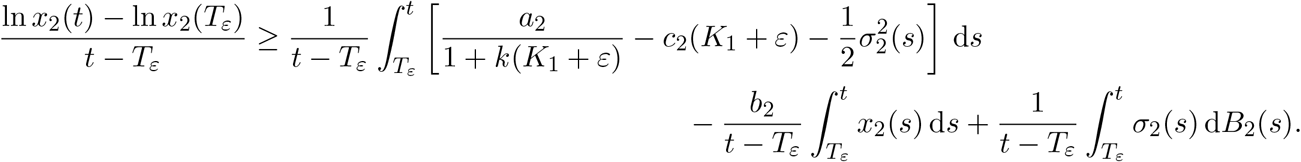

On using the law of strong number and by the standard persistence lemma [40, 53],

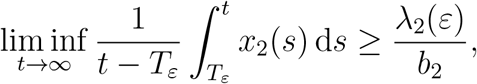

where 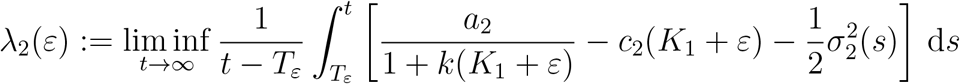.

Since *T*_*ε*_ is a finite real number, on letting *ε* → 0^+^, we have *λ*_2_(*ε*) → *λ*_2_. Hence

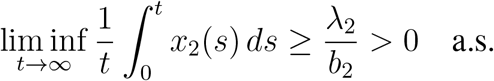

